# Butterflyfishes as a System for Investigating Pair Bonding

**DOI:** 10.1101/214544

**Authors:** Jessica P. Nowicki, Lauren A. O’Connell, Peter F. Cowman, Stefan P. W. Walker, Darren J. Coker, Morgan S. Pratchett

## Abstract

For many animals, affiliative relationships such as pair bonds form the foundation of society, and are highly adaptive. Animal systems amenable for comparatively studying pair bonding are important for identifying underlying biological mechanisms, but mostly exist in mammals. Better establishing fish systems will enable comparison of pair bonding mechanisms across taxonomically distant lineages that may reveal general underlying principles. We examined the utility of wild butterflyfishes (f: Chaetodontidae; g: *Chaetodon*) for comparatively studying pair bonding. Stochastic character mapping inferred that within the family, pairing is ancestral, with at least seven independent transitions to group formation and seven transition to solitary behavior from the late Miocene to recent. In six sympatric and wide-spread species representing a clade with one ancestrally reconstructed transition from paired to solitary grouping, we then verified social systems at Lizard Island, Australia. *In situ* observations confirmed that *Chaetodon baronessa, C. lunulatus*, and *C. vagabundus* are predominantly pair bonding, whereas *C. rainfordi, C. plebeius*, and *C. trifascialis* are predominantly solitary. Even in the predominantly pair bonding species, *C. lunulatus*, a proportion of adults (15 %) are solitary. Importantly, inter- and intra-specific differences in social systems do not co-vary with other previously established attributes (geographic occurrence, parental care, diet, or territoriality). Hence, the proposed butterflyfish populations are promising for comparative analyses of pair bonding and its mechanistic underpinnings. Avenues for further developing the system are proposed, including determining whether the utility of these species applies across their geographic disruptions.

## Introduction

Social bonds are foundational to many animal societies [1, 2]. Arguably, the most extreme form of social bond is the pair bond—a selective, relatively pro-social and enduring affiliation between 2 individuals that is maintained beyond (or outside of) reproduction. Pair bonding has independently evolved numerous times within and across major vertebrate lineages [3–7], where it represents a defining feature of species-typical social structure [8], and can shape aspects of parental [9–11] and mating [12, 13] strategies. The adaptive significance of vertebrate pair bonding is relatively well understood (reviewed by [3–5, 7, 11, 14–16]). Research is increasingly focussed on the mechanistic (e.g., neuroendocrine) basis of pair bonding, largely due to its implications for the biological mechanisms of human pro-sociality, anti-social psychological disorders [17, 18], and physical health [19]. However, most of what is known about the mechanistic basis of pair bonding comes from extensive studies on a single genus, *Microtus* voles (reviewed in [20–27]). A scarcity of complementary research among other organisms, has led to little being known about the mechanistic basis for the evolution of pair bonding across vertebrates, which ultimately makes it difficult to identify general principles for the sub-phylum. Moreover, because current animal systems for comparatively studying pair bonding are confounded with other life-history attributes (e.g., also display parental care), it is difficult to identify causal mechanisms of pair bonding apart from other behavioral correlates. Expanding pair bonding systems to better include teleost fishes is a promising solution to these limitations, owing to their distant taxonomic relation to mammalian and avian systems [28], unparalleled species diversity [29], and extreme diversity in social systems, ecology, and behaviour [30, 31].

*In situ* behavioral observations on wild organisms are a critical first step towards establishing the existence and variation of social systems within and among species [32–35]. Species that exhibit inter-individual variation in social systems are particularly useful for comparatively identifying mechanistic correlates [36, 37] when potential confounds such as geographic occurrence, life history, and behavioral ecology are controlled. Whereas, inter-species comparisons within a taxon; when controlling for aforementioned confounds and phylogenetic relatedness; can inform mechanistic correlates related to how social systems have evolved across phylogenetic time [38–40], potentially illuminating general principles for the taxon that may not be apparent in a single species [41, 42]. While systems for comparatively studying the mechanisms of pair bonding were originally limited to a single genus of mammal, *Microtus* voles [43, 44], additional comparative systems for other taxon within mammals and other major lineages have recently emerged: mammals: *Peromyscus* mice [45]; birds: *Coturnix* quails [46]; teleosts: *Neolamprologus, Telmatochromis* [47] and *Herichthys* [48] cichlids. A challenge among all established comparative pair bonding systems is that in one system or another, species differences in pairing phenotype co-vary with differences in other attributes, including parental care, general social affiliation, and territoriality [46–53]. Since shared neuroendocrine mechanisms have been shown to regulate all of these attributes [8, 54], it is difficult to use these systems to isolate independent causal mechanisms of pair bonding without additional experimental validation being required.

Teleost fishes offer many opportunities for comparatively studying social systems [31, 55]. Among vertebrates, the lineage is the most taxonomically diverse (~29,000 described species) [29] and displays extreme variation of social behavior among individuals and species [30, 31]. The family Chaetodontidae (butterflyfishes and bannerfishes; “chaetodontids”) is attractive for comparative research into pair bonding specifically. Chaetodontidae are widely distributed throughout the world’s oceans, occurring in all coral reef regions [56, 57]. The family includes at least 127 extant species [56], 77 of which predominantly occur in paired social groups (data sourced from [57–64]), ostensibly accounting for ~21% of all reported pair bonding marine fishes (data sourced from [5, 7]). Their evolutionary history is also relatively well understood with about 75% of the family represented in dated molecular phylogeny [65, 66]. *Chaetodon* butterflyfishes have undergone rapid species diversification relatively recently (~ 16 Ma) [66], resulting in 93 nominal extant species, among which the majority (59 spp.) predominantly occur in paired social groups (data sourced from [57–64]). Available data on select pairing species suggests that pairs exhibit partner fidelity. Partners have been shown to remain together for the full duration of monitoring studies, which range from several months to 7 years [58, 67–71]. Such duration of partner fidelity can be considered prolonged to long-term, since *Chaetodon* spp. consistently live for more than 10 years [72]. By contrast, a minority of *Chaetodon* spp. predominantly occur in solitude or gregarious (3+ individual) groups [58, 73, 74], suggestive of species diversity in social systems. As species diversity in Chaetodontid behavioural ecology [60, 75, 76], biogeography [77, 78], and relatedness [66, 79, 80] is well established, comparisons of social systems can be made in a highly controlled manor. To that end, all chaetodontids are broadcast spawners that effectively display no parental care [58, 81], and would therefore provide the first insights into the mechanistic correlates of pair bonding independent of parental care.

Although there are numerous studies on Chaetodontidae social behaviour, surprisingly few studies have established species’ typical social systems [60], defined by a whole of interactions and relationships between individuals, such as social grouping, aggression, social bonding, group sex composition and mating systems [82]. Consequently, controlled comparative systems for studying variation in pair bonding are yet to be developed for the group. Additionally, because the evolutionary history of pair bonding within Chaetodontidae remains unexamined, such comparative systems are yet to be developed within a framework that considers ancestral states.

The aim of this study was to develop a new *Chaetodon* butterflyfish system for controlled and evolutionarily-informed intra- and inter-species comparative studies on pair bonding. Specifically, we first sought to conduct ancestral reconstruction analysis to inform the evolutionary history of chaetodontid sociality. Following, for 6 species, we sought to verify intra- and inter-species variation in social systems (i.e., pair bonding vs. solitary living) through *in situ* field studies on wild populations. To do so, we focused on features that are routinely recognized as characteristic of pair bonding across taxa, that are useful for distinguishing pair bonding from non-pair bonding phenotypes, and that are ecologically relevant to butterflyfishes. These features include i) the predominance of group sizes of 2 individuals [74, 83–86], ii) selective affiliation with a distinct partner [22, 83, 84], which in the case of fishes may be expressed as proximate and parallel (i.e., “pair”) swimming [58, 87], iii) selective aggression towards non-partners [24, 35, 46, 67], iv) predominantly heterosexual pair composition [14, 32, 58, 87–89], and v) long-term partner fidelity/endurance [14, 58, 70, 71, 83–85, 90, 91]. Specifically, we tested the predictions that 3 species would predominantly occur in heterosexual, enduring pairs that exhibit selective affiliation towards partners over non-partners, and selective agonism towards non-partners over partners; whereas 3 would predominantly occur in solitude, and exhibit infrequent and indiscriminate affiliation with another individual. Finally, the same predictions were tested among individuals of 1 species, *Chaetodon lunulatus*.

## Materials and Methods

### Evolutionary history of Chaetodontidae sociality

The most completely sampled and date phylogeny was chosen to conduct ancestral reconstruction of social group sizes in the family Chaetodontidae. The phylogeny of [66] includes 95 of the 127 described species and is based on four mitochondrial and four nuclear genes. Briefly, the 8-gene dataset was exposed to Maximum Likelihood analysis in the program Garli [92], with the best ML topology chosen as a starting tree for Bayesian age estimation analyses with fossil calibrations in the program BEAST [93]. This resulted in a posterior distribution of dated trees, which were then summarize as a maximum clade credibility tree (MCC).

A literature search was conducted to classify the predominant social group size of all chaetodontid species as either ‘pairing’, ‘gregarious’ (forming groups of 3 or more) or ‘solitary’. From this literature search, 79 species were classified as pairing, 17 were classified as gregarious and 14 were classified as solitary (Table S1). Of the remaining species, 1 species has been recorded as displaying both pairing and gregarious behavior (*Chaetodon gardineri*, although this species was not sampled in the phylogeny) and the sociality of 17 species could not be determined. Overall, there were 20 species with group size data that were missing from the phylogeny. Species sampled in the phylogeny where no accurate determination could be made on group size (*Amphichaetodon melba*, *Chaetodon blackburnii, Prognathodes marcellae, P. aya*) were coded as having an equal probability of being in any of the 3 states, allowing their probable state to be reconstructed during the ancestral reconstruction analyses.

The evolutionary history of Chaetodontidae social grouping was explored using a stochastic character mapping function from the R package phytools [94]. The stochastic character mapping procedure samples simulated histories of a trait across the evolutionary history of a phylogeny, and can incorporate topological uncertainty by conducting the analyses across a distribution of trees. Using this method, we examined transition rates among sociality character states and highlight the temporal origins of group sizes. To begin, we ran 1000 stochastic character maps on the MCC tree of Cowman and Bellwood [66] using the ‘make.simmap’ function of phytools with Q = “mcmc” to sample the transition matrix (Q) from its posterior probability distribution. The mean transition matrix from this analysis was then used to infer the stochastic character mapping of 1000 tree topologies sampled from the posterior distribution of tree taken from the Cowman and Bellwood [66] study. For each tree, 10 stochastic maps were generated resulting in 10,000 mappings. From this set of stochastic character maps, the average number of transitions among character states were calculated, and character histories were summarized as state probabilities on the internal nodes of the MCC tree.

### Species and study site

*Chaetodon lunulatus, C. baronessa, C. vagabundus, C. plebeius, C. rainfordi*, and *C. trifascialis* were selected as candidates for establishing designs for examining intra- and inter-species variation in pair bonding, for several reasons. Firstly, available evidence suggests that these species may exhibit dichotomous social systems, with *C. baronessa, C. lunulatus*, and *C. vagabundus* appearing to be predominantly pair bonding, and *C. rainfordi, C. plebeius*, and *C. trifascialis* appearing to be primarily solitary [32, 60, 73, 74, 95]. Apart from *C. plebeius*, this apparent species diversity in social systems appears to be highly consistent throughout their geographic distributions (data sourced from [32, 58, 60, 73, 74, 89]). However, as with most chaetodontids, the social systems of these species at a given geographic location are often inferred from few social proxies (mostly predominant group size) (e.g., [60, 74, 89, 95]) rather than verified by quantitatively and holistically assessing a repertoire of social behaviors that cumulatively define social systems (*C. lunulatus* (= *trifasciatus*) at Yaeyama Islands notwithstanding [67, 68, 74, 96]). Hence, reliable assessments of social systems of these species (*C. lunulatus* notwithstanding once more) and among these species at a single location remain absent. Secondly, these species are closely related congeners [66, 80] that are widely distributed throughout the Indo-/Western-Pacific region [29], wherein they can be found in relative abundance and co-occurring in sympatry [32, 74, 97]. Finally, the co-occurring populations at Lizard Island, located in northern section of Australia’s Great Barrier Reef (14°40’S, 145°27’E), were chosen for this study, because their feeding ecology [97–100], territoriality [101], demography [72], and habitat associations [98] have been previously established; and importantly, do not co-vary with predicted social systems. All field studies were conducted on the north-western side of the island, where there are numerous distinct platform reefs that are easily accessible. Only individuals that were at least 80% of average species-specific asymptotic body length, and therefore likely reproductively mature [89] were considered. Studies were conducted at haphazard times from 0800-1800 hrs, from January – May, 2013-2015.

### Verifying inter- and intra-specific variation in social systems

#### Determining species-predominant group size

Social systems were first assessed by determining species’ predominant group sizes. For each species, group size frequencies were measured at 5 haphazardly selected reefs using 6 replicate 50 X4 m belt transects per reef. During surveys, each individual (or group of individuals) within the transect area was followed for a 5 min observation period. Group size was determined by the number of individuals (either 1, 2, or 3+ individuals) that displayed proximate swimming (within 1.5 m distance) for at least 3 consecutive min during the 5 min observation period. Swimming distance was visually estimated after practicing accuracy on dummy fishes prior to the study. Sample sizes of observations varied in accordance with variation in abundance: *C. rainfordi* (n = 48), *C. plebeius* (n = 61), *C. baronessa* (n = 76), *C. lunulatus* (n = 98), *C. trifascialis* (n = 43), *C. vagabundus* (n = 55). To determine species’ predominant group size, for each species, the total number of observations of different group sizes were pooled across reef sites, and compared to a pre-defined uniform distribution that would be expected if individuals had no preference for any group size (33.33% of observations in each group size) using a chi square goodness-of-fit.

### Within- and between-group agonism and affiliation

To further explore social systems, field observations were conducted to measure social affiliation and agonism within and between conspecific groups. *In situ* behavioural observations were conducted on snorkel across 5 haphazardly selected reefs. Focal individual(s) within the group were identified, and then observed from a distance of 2-3 metres. Focal individuals were allowed 3 min to acclimate to observers’ presence. Time spent proximate swimming, defined as swimming within a 1.5 m distance from another conspecific; and parallel swimming, defined as swimming faced within a 315-45° angle relative to the faced position of another conspecific (Fig 1), were sampled once every 10 sec throughout a 3 min observation. Swimming distance and angle were estimated visually. For predominantly paired species, these behaviours were measured towards both partner and non-partner conspecifics; for predominantly solitary species and solitary *C. lunulatus*, they were measured towards other conspecifics. While we attempted to sample both proximate and parallel swimming for each fish observed, there were few cases in which only one of these behaviours were measured. Sample sizes of observations for each of proximate and parallel swimming behaviours are as follows: *C. rainfordi* (n=14, each behaviour), *C. plebeius* (n=15, each behaviour), *C. baronessa* (n=18 and n=20, respectively), paired *C. lunulatus* (n=18, each behaviour), solitary *C. lunulatus* (n=16, each behaviour), *C. trifascialis* (n=15, each behaviour), and *C. vagabundus* (n= 24 and 17, respectively). Because it was determined in the present study that *C. lunulatus* is both pairing and solitary, for this species, proximate and parallel swimming with another conspecific was compared between paired and solitary individuals using non-parametric Mann-Whitney *U*-test (SPSS Software), due a lack of normality of residual variance. For predominantly paired species, total agonistic acts, including staring, chasing, fleeing, and encircling (see [67] for detailed description) towards partners and non-partner conspecifics, were measured. Sample sizes for observations of agonistic acts were as follows: *C. baronessa* (n= 26), *C. lunulatus* (n= 25), *C. vagabundus* (n= 24).

**Fig 1.**
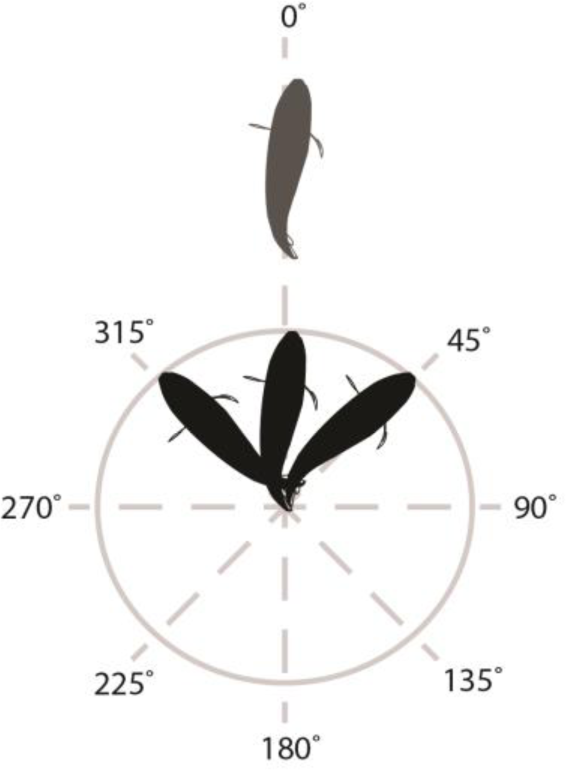
Schematic of parallel swimming examined in six *Chaetodon* species. Parallel swimming by the focal fish (black) was defined as being faced within a 315-45° angle relative to the faced position of the conspecific (grey), whose faced position was designated 0°.

### Sex composition

To examine the sex composition of pairs among predominantly pairing species, a sub-sample of pairs were collected following behavioural observations and sacrificed. Individuals of predominantly solitary species and solitary *C. lulnulatus* were also collected for sex composition analysis. Gonads were removed and fixed in formaldehyde-acetic acid-calcium chloride solution (FACC) for at least 1 week. Thereafter, gonads were dehydrated in a graded alcohol series, washed in xylene, embedded in paraplast, sectioned transversely (7 µM thick), and stained with hematoxylin and eosin. Sections were examined under a compound microscope (400 X magnification) for the presence of characteristic sex cells [102, 103]. Among pairs, 3 categories of sex composition were found: heterosexual pairs, homosexual pairs, and pairs in which at least one individual was ostensibly a hermaphrodite. To statistically test whether paired individuals had a preference for partnerships of a particular sex composition, the number of pairs in different pair sex composition categories was compared to a pre-defined uniform distribution that would be expected if individuals had no preference for a given pair sex composition (33.33% of all pairs in each category) using a chi-squared goodness-of-fit.

### Partner fidelity in pairs

To test whether the 3 predominantly pairing species of this study exhibit partner fidelity, we uniquely tagged pairs of each species (*C. baronessa*, n=12; *C. lunulatus*, n= 18; *vagabundus*, n= 17) and then re-surveyed tagged fishes after 6-weeks to record changes in partner identity (i.e., pair permutation). Paired fishes were identified as described above, and then caught using a barrier net. Paired individuals were tagged on opposite sides of the dorsal musculature with unique and matching colour coded external tags using a hand-held tagging applicator (Floy T-bar Anchor) [104]. Tagged individuals were re-assessed for partner fidelity after 6 weeks, as this duration would inform the extent of short-term partner fidelity. A team of 3 snorkelers used an "expanding circle" search approach to reidentify tagged butterflyfishes. Once tagged fishes were detected, 3 min observations at a distance of at least 2 m from fish were again conducted to test for partner affiliation (as above); and respective partners were then carefully examined at within 1 m to determine identity (i.e., tagged and known/untagged and unknown). To facilitate re-detection of tagged fishes, this study was conducted on a single distinct platform reef, separated from nearby reefs by an open expanse of sand, which was expected to minimise movement of fishes away from the vicinity in which they were originally tagged. We had also planned to assess partner fidelity over several years; however, were unsuccessful due to tags having been dislodged from fishes by the next date of re-assessment (11 months post tagging), made apparent by no tagged fishes being re-identified within the study site, and yet scars observed on the tagging location of several fishes within the study site.

## Results

### Evolutionary history of Chaetodontidae sociality

Results from the 10,000 stochastic character mappings summarized on an MCC tree (figure 2) identify pair bonding as the most likely ancestral sociality of the family. Several independent transitions were recorded from pair bonding to solitary (average of 7.5 transitions) and gregarious behaviour (average of 7.1 transitions; Fig 2 inset). Reversions back to pair bonding from gregarious or solitary lineages appear to be uncommon. While a subclade with the *Chaetodon* Clade 3 (CH3) appears to retain the transition to solitary behaviour for much of its evolutionary history (with some changes to gregarious and pair forming), there was very little diversification observed within lineages reconstructed as displaying gregarious grouping. Except for the expansion of the *Hemitaurichthys* lineage in the Bannerfish clade, gregarious behaviour is only reconstructed along extant, recent lineages across the great *Chaetodon* clade (CH2, CH3, CH4). As for the species that were coded as having equal probabilities of being in either of the three states, based on their position in the phylogeny and the reconstruction, both *Proganthodes* species are reconstructed and mostly solitary, while both *Amphichaetodon melbae* and *Chaetodon blackburnii* are reconstructed as most likely pair bonding (Fig 2).

**Fig 2.**
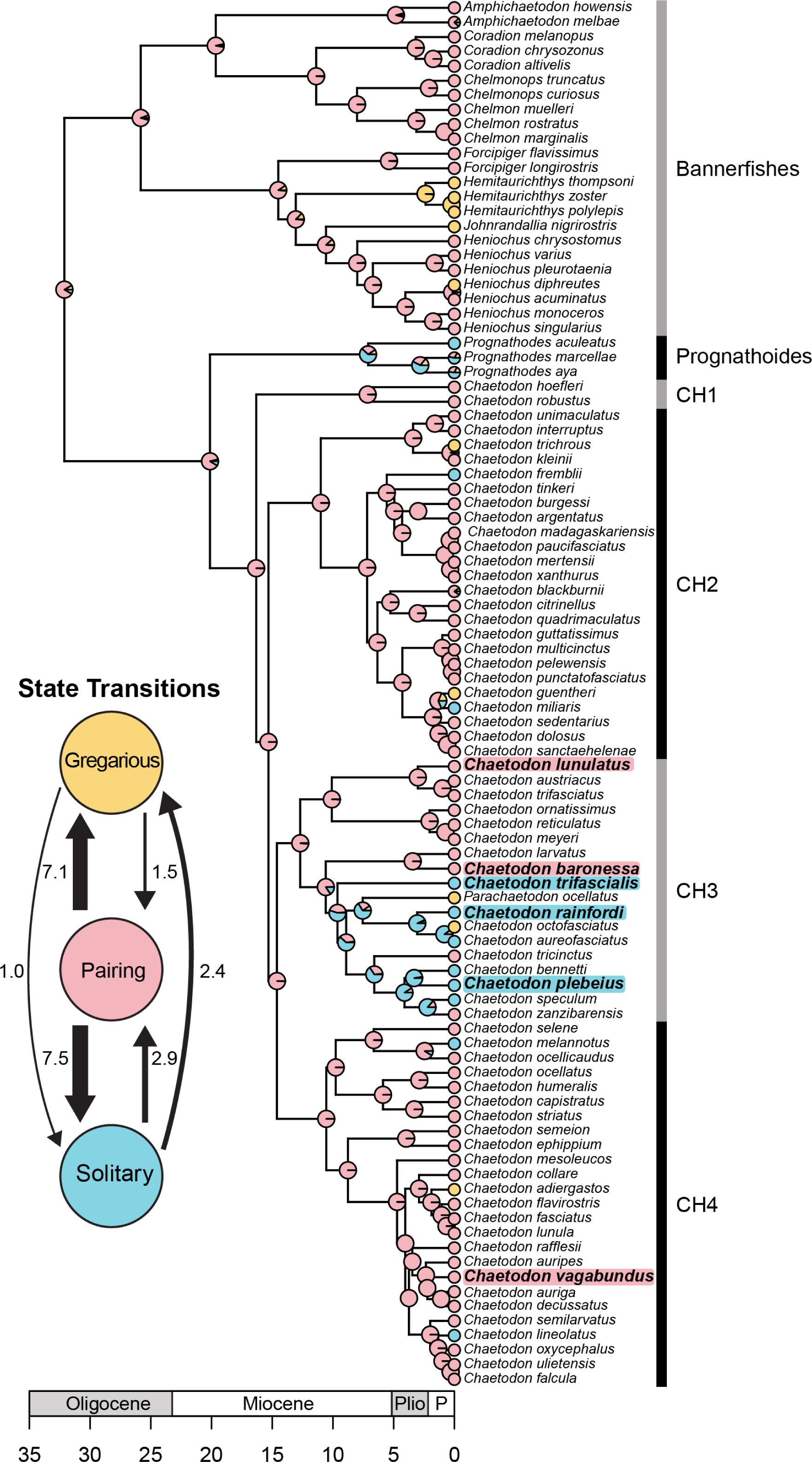
Ancestral reconstruction of social behavior in the family Chaetodontidae summarized on a published maximum clade credibility tree ([66]). Pie charts at nodes represent the posterior probabilities of state reconstructions, summarized from 10,000 stochastic character maps across 1000 randomly sampled topologies from the BEAST posterior distribution of trees [66]. Within the family, pairing is reconstructed as the ancestral character with several subsequent independent transition to solitary and gregarious grouping with few reversals to pairing (inset). Within the study group (highlighted in blue for solitary and pink for pairing), pairing is an ancestral state, with 1 potential origins of solitary sociality in the common ancestor to C. trifascialis, C rainfordi, and C. plebeius.

### Inter- and intra-specific variation in social systems

#### Species predominant group size

For all 6 species, the distribution of observations of groups across different group sizes differed significantly from a uniform distribution (*C. baronessa*: χ^2^ = 73, df = 2, *p* <0.001; *C. lunulatus*: χ^2^ = 114, df = 2, *p* < 0.001; *C. vagabundus*: χ^2^ = 42, df = 2, *p* < 0.001; *C. rainfordi*: χ^2^= 64, df = 2, *p* < 0.001; *C. plebeius*: χ^2^ = 89, df = 2, *p* < 0.001; *C. trifascialis*: χ^2^ = 41, df = 2, *p* < 0.001). There was also an apparent dichotomy in predominant group size across species. Regardless of study site, *C. baronessa, C. lunulatus*, and *C. vagabundus* had a predominant group size of 2 individuals, with 78, 84, and 71 % of individuals found in pairs, respectively; and were seldom found in a group size of 1 individual (22, 15, and 29 % of observations, respectively) (Fig 3). Among predominantly pairing species, group sizes of 3+ were only ever observed for *C. lunulatus*; however, this was only on one occasion. By contrast, *C. rainfordi, C. plebeius*, and *C. trifascialis* had a predominant group size of 1 individual (88, 90 and 80 %, respectively) (Fig 3). Individuals of these species were less commonly observed paired (10-15%), and very rarely observed in a group size of 3+ (1-2 %).

**Fig 3.**
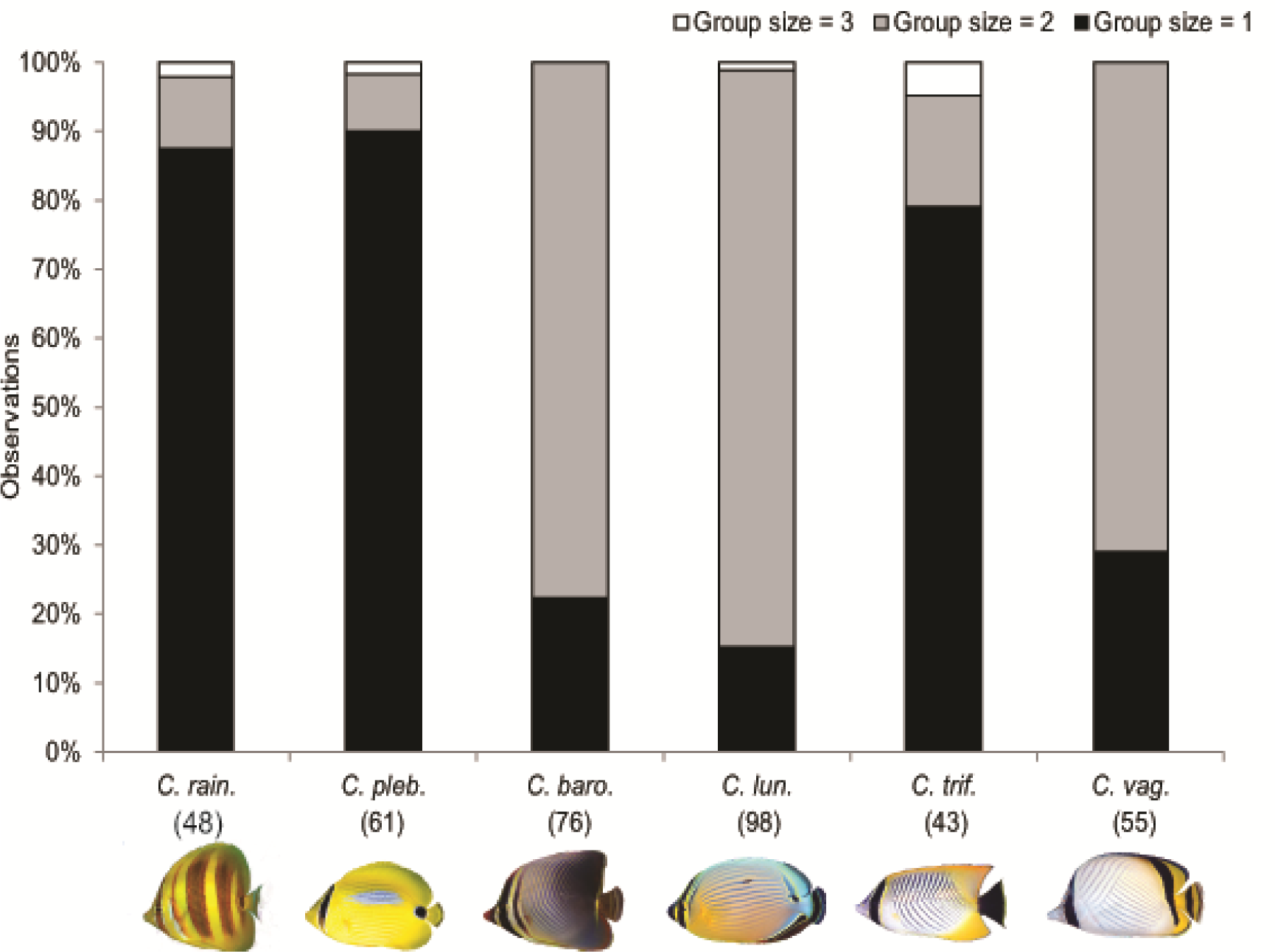
Group size frequency distribution of six *Chaetodon* spp. at Lizard Island. Numbers in parentheses indicate sample size of total observations of groups.

### Level of selective proximate and parallel swimming

The occurrence of proximate and parallel swimming clearly distinguished paired versus solitary grouped species (Fig 4A, B). Pairs of *C. baronessa, C*. *lunulatus* and *C. vagabundus* ranged as a single coordinated social unit throughout the reef, spending the majority of time (72 ± 7.41, 89 ± 6.2, and 81 % ± 6.1 SE, respectively) swimming within 1.5m of their partner, and most of the time (53 ± 8.1, 72 ± 5.8 SE, and 69 % ± 6.6 SE, respectively) were faced within a 315-45° angle of their partner (see Fig 1). By contrast, singletons of *C. rainfordi, C. plebeius*, and *C. trifascialis* displayed no apparent social affiliation with another individual, as they spent 100 % of their time swimming further than 1.5 m from another conspecific; and most commonly, no other conspecific was within a field of view. Similarly, proximate and parallel swimming significantly and strongly varied between paired and solitary grouped *C. lunulatus* individuals (proximate swimming: *U* = 9, *p* < 0.001; parallel swimming: *U* = 9.5, *p* < 0.001) (Fig 4A, B). While paired individuals displayed these behaviors exclusively with their partners and at relatively high levels (swimming within 1.5 m from partner for 89 % ± 6.2 SE of time; swimming faced within a 315-45° angel of their partner 72 % ± 5.8 SE of the time), solitary individuals displayed these behaviors at relatively low levels (swimming within 1.5 m from another conspecific 3.1 % ± 2.3 of time; swimming faced within a 315-45° angle of another conspecific 2.8 % ± 1.5 of the time).

**Fig 4.**
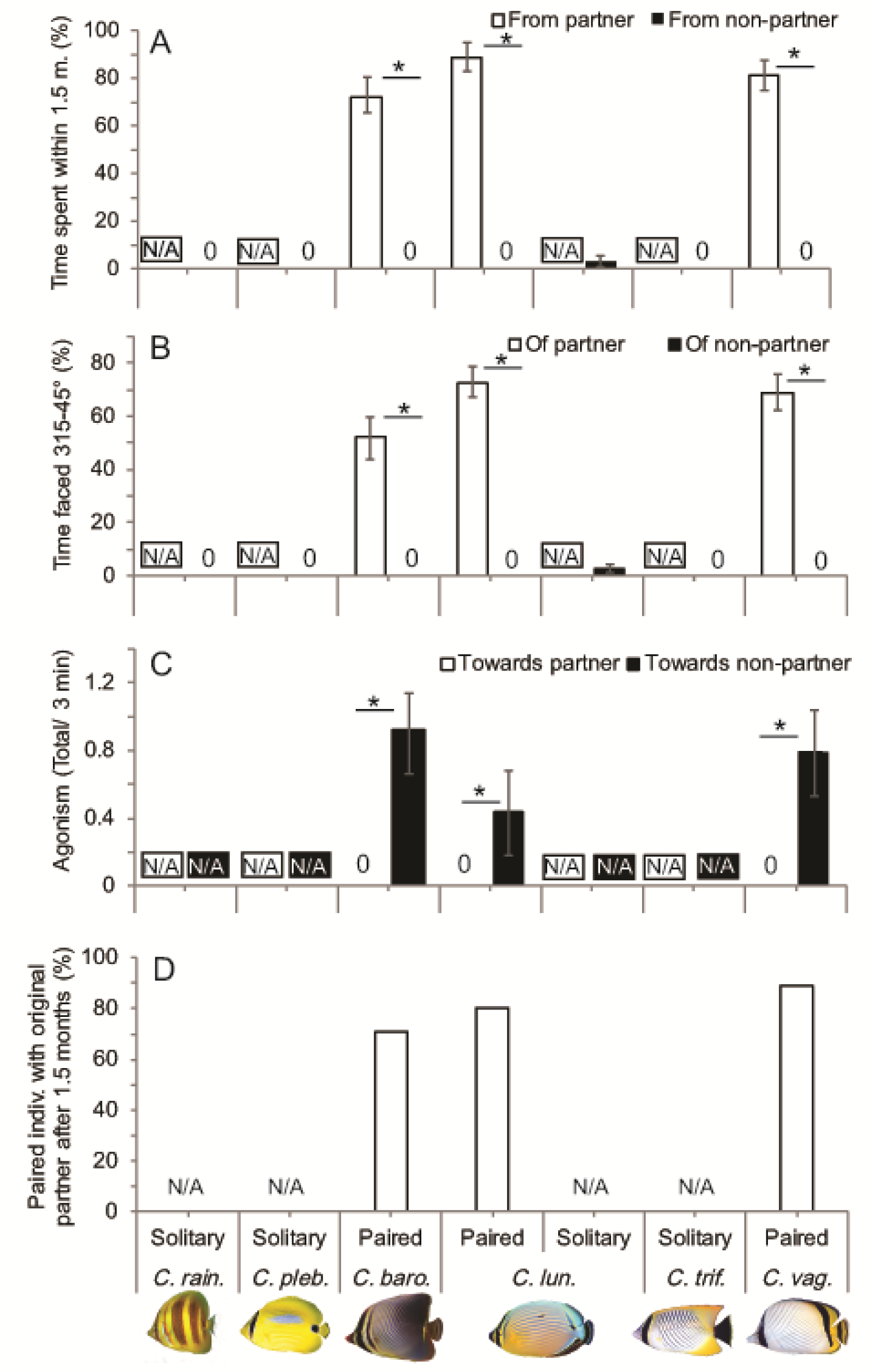
Differences in social behaviors between predominantly paired and solitary grouped *Chaetodon* spp. and *C. lunulatus* individuals. (A) Mean % ± SE time spent proximate swimming with another conspecific. (B) Mean % ± SE time spent parallel swimming with another conspecific. (C) Mean agonism ± SE towards partner vs. non-partner conspecifics among pairs. (D) Percentage of pairs displaying partner fidelity after 6 weeks. Asterisks indicate significant differences between groups.

### Level of selective agonism of pairs

Pairs of *C. baronessa, C. lunulatus*, and *C. vagabundus* displayed agonism exclusively towards non-partner conspecifics (Fig 4C), and no agonism towards partners. However, even agonism towards non-partner conspecifics was infrequent and minor, consisting mostly of staring displays.

### Partner fidelity in pairs

Across the 3 pairing species, a total of 26 of the original 94 tagged fish were re-identified after 6 weeks: *C. baronessa*: 7/24 fish, *C. lunulatus*: 10/36 fish; *C. vagabundus*: 9/ 34 fish, and within their original general reef location. Among these re-identified tagged fish, individuals were paired with their original partners in 5/7 (71% of) cases for *C. baronessa*, 8/10 (80% of) cases for *C. lunulatus*, and 8/9 (89% of) cases for *C. vagabundus* (Fig 4D); were paired with a different (non-tagged) fish in 1/10 (10 % of) cases for *C. lunulatus*, and 1/9 (11% of) cases for *C. vagabundus*; and were found solitary in 2/10 (20% of) cases for *C. baronessa*, and in 1/9 (11% of) cases for *C. lunulatus*; In cases where re-identified focal fish were not found with their original partners, their original partners were not found by the observers.

### Sex composition

Among predominantly pairing species, most of the pairs for which we determined sex histologically were heterosexual, whereas homosexual pairs or pairs comprised of at least one ostensive hermaphrodite were uncommon (Table 1). The frequency of heterosexual pairs differed significantly from a uniform distribution (*C. baronessa*: *X*^2^ = 17.7, df = 2, *p* < 0.001*; C. lunulatus*: *X*^2^ = 19.2, df = 2, *p* < 0.001; *C. vagabundus*: *X*^2^ = 12.0, df = 2, *p* < 0.001). Among predominantly solitary species, singletons were mostly female and uncommonly male or ostensive hermaphrodites (Table 2). Among solitary *C. lunulatus*, singletons were equally male and female (Table 2).

**Table 1.** Predominantly pairing species in current study: sex composition of pairs

| Species | Hetero. pairs (n) | Homo. pairs (n) | Hermaph.* pairs (n) | Total pairs | Hetero. pair ratio | Homo. pair ratio | Hermaph. pair ratio |
| --- | --- | --- | --- | --- | --- | --- | --- |
| *C. baronessa* | 12 | 2 (F + F) | 0 | 14 | 0.86 | 0.14 | 0.00 |
| *C. lunulatus* | 13 | 1 (M + M) | 1 (F + H) | 15 | 0.87 | 0.07 | 0.07 |
| *C. vagabundus* | 6 | 0 | 0 | 6 | 1.00 | 0.00 | 0.00 |

Key: F = female; Hetero. = heterosexual; H/Hermaph. = ostensive hermaphrodite; Homo. = homosexual; M = male. \*Pairs for which at least one partner was a hermaphrodite.

**Table 2.** Predominantly solitary species and solitary *C. lunulatus* in current study: sex composition of individuals

| Species | Males (n) | Females (n) | Hermaph. (n) | Total indiv. | Male ratio | Female ratio | Hermaph. ratio |
| --- | --- | --- | --- | --- | --- | --- | --- |
| *C. rainfordi* | 2 | 12 | 1 | 15 | 0.13 | 0.80 | 0.07 |
| *C. plebeius* | 0 | 14 | 1 | 15 | 0.00 | 0.93 | 0.07 |
| *C. trifascialis* | 2 | 12 | 1 | 15 | 0.13 | 0.80 | 0.07 |
| *C. lunulatus* | 6 | 5 | 0 | 11 | 0.55 | 0.45 | 0.00 |

Key: Hermaph. = ostensive hermaphrodite.

## Discussion

### Evolutionary history of Chaetodontidae sociality

This is the first study of the evolutionary history of Chaetodontidae sociality. Stochastic character mapping on 1000 posterior trees topologies (10 maps per tree) revealed that pairing appears ancestral and moderately conserved (Fig 2), with transitions to solitary and gregarious grouping occurring only in the last 10-15 million years. Only in one instance did the transition to solitary behavior have any subsequence diversification within a Chaetodon subclade (CH3 Fig 2). This transition occurred in a common ancestor of three of our study species that reside within clade CH3 (*C. rainfordi*, *C. plebeius*, and *C. trifascialis*). This clade also includes potential transitions to gregarious behavior (Parachaetodon ocellatus, and C. octofasciatus) and two independent reversions to pair bonding behavior (C. tricintus, C. zanzibarensis). It is likely that the inclusion of unsampled Chaetodon species in future phylogenies might alter the character reconstructions highlighted here, but based on the current systematic of the family, only 3 species are missing from clade CH3 (*C. triangulum, C. melapterus*, and *C. andamanensis*), all of which were found to be pairing. *C. andamanensis* is the only likely species to be placed within the CH3 subclade representing a transition to solitary behavior (most probably as a sister species to *C. plebeius*), potentially representing another reversion to pair bonding within the clade.

### Intra- and inter-specific variation in social systems of Chaetodon butterflyfishes

This is one of the few studies to formally characterize the diversity of social systems within and among several butterflyfish species, including those inhabiting the same geographic location (see also [32, 105]). Results support our initial hypothesis that at Lizard Island, *C. baronessa, C. lunulatus*, and *C. vagabundus* are predominantly pair bonding, while *C. rainfordi, C. plebeius*, and *C. trifascialis* are predominantly solitary. They moreover meet our expectation that in one species, *C. lunulatus¸* both pair bonding and solitary living occurs among individuals. This reaffirms that butterflyfishes exhibit considerable diversity in social systems—an assumption that has been largely based on sparse behavioral observations, and primarily predominant group size [60, 74, 95].

### Intra-specific variation in *Chaetodon lunulatus*

We found that at Lizard Island, *C*. *lunulatus* occurs in pairs 90 % of the time. Heterosexual pairing predominates, occurring significantly more often than expected by chance alone (87 % of the time), and therefore appears to be selected for. Pairs display a high level of proximate and parallel swimming (89 % and 73 % of time, respectfully) that occurs exclusively between partners. Even when partners were not swimming “proximately” (≤ 1.5 m), they almost always remain within close range (≤ ~4 m) of each other. While agonism in paired individuals is infrequent, it occurs exclusively towards non-partner conspecifics. Among the 9 pairs that were re-identified, 8 persisted with their original partners throughout the duration of the study (6 weeks). While we had hoped to measure partner fidelity for much longer time period (> 12 months), this was not feasible, due to loss of tags after 6 weeks. (In future studies, we suggest longer-term assessment of partner fidelity using unique markings on focal individuals for identification [68, 90] in preference to external tags.) When taken together, these observations verify that *C. lunulaus* is predominantly and strongly pair bonding at Lizard Island.

Consistent with Lizard Island, other populations of *C. lunulatus* invariably display a predominant social group size of 2 individuals, and indeed exhibit the highest prevalence of paired grouping among butterflyfishes overall [60]. Ninety-five percent of observations at Yaeyama Island (Japan) [74], 81 % at Moorea Island (French Polynesia) [73], 84 % at Heron Island (Australia) [32], 76 % at Marshall Islands (Australia) [32], and 68 % at Palm Island (Australia) [89] are of paired groups. Reese (1975) reported a relatively low pairing ratio (53%) at Johnston Island (Hawaii); however, this was from a relatively low sample size (n=17 total observations). Adult pairs are predominantly heterosexual (92 %), presumably in order to facilitate reproduction [89]. Among pairs, partners display highly affiliative pair swimming, maintaining coordination and close proximity whilst ranging throughout the reef [32], and particularly a shared long-term territory [90, 96], likely functioning as a form of territory defense that conspicuously advertise occupancy [58]. Within the Yaeyama Islands (Japan) population, for example, partners spend 89 % of their time swimming within 2 m of each other, and only 11 % of their time swimming at further distances [96]. Aggression between partners rarely occurs, and may be consequent of failed partner recognition [96]. By contrast, aggression towards non-partners is well documented in the species [32, 67, 95, 101, 106, 107], including the Lizard Island population [95, 101, 107], where it is attributed to territory defense [32, 101, 106, 107] and mate-guarding [67]. However, territorial aggression in *C. lunulatus* pairs is generally passive [74, 106], consistent with the ‘dear enemy’ model of low-cost resource defense once territories have been established among neighbors [14, 108]. Partner fidelity in *C. lunulatus* has been previously examined only once, where individuals remained paired with the same partner for up to 7 years (Heron Island, Australia) [69]. Finally, partnerships are predominantly (93 % of pairs, Palm Islands, Australia) [89], if not exclusively (100 % of pairs, Heron Island, Australia) [32] heterosexual; and in one study it was shown that mating occurs exclusively within the pair (Yaeyama Islands, Japan) [68]. Hence, *C. lunulatus* is predominantly and strongly pairing throughout its geographic range, and findings from specific populations suggest that these partnerships are both socially and reproductively monogamous.

At Lizard Island, we recorded that 15 % of *C. lunulatus* adults occur in a group size of 1 individual that rarely exhibits proximate or paralleled swimming with another conspecific (~3% of time); and hence, are solitary. It is possible that in certain cases, paired individuals were mistaken for singletons. However, this would have occurred infrequently, since among pairs, partners spend nearly all their time swimming within 4 m of each other; and yet in nearly all instances where individuals were recorded as solitary, another conspecific was not within field of view. Similarly, among the Yaeyama Islands (Japan), Moorea Island (French Polynesia), Heron Island, Marshall Island (Australia), and Johnston Island (Hawaii) populations; solitary groups occur on average 7 % of the time. Elsewhere (Johnston Island, Hawaii), the proportion of solitary individuals is as high as 47 % [32]. In any population, a small proportion of mature individuals would be expected to be solitary due to partner scarcity or loss [109]. It is also possible that there are differences in the propensity to pair bond vs. remain single within and between populations, due to differences in selective pressures (e.g., food competition).

### Inter-specific variation among *Chaetodons*

We found that at Lizard Island, as in *C. lunulatus; C. baronessa* and *C. vagabundus* are predominantly found in pairs (78 and 71 % of observations, respectively), infrequently in solitary groups (22 % and 29 % of observations, respectively), and never in gregarious groups. Pairs are predominantly if not exclusively heterosexual (*C. baronessa*: 86 % of pairs; *C. vagabundus*: 100 % of pairs). The occurrence of heterosexual pairing in these species is significantly higher than that expected by chance alone, indicative of positive selection. Paired individuals of *C. baronessa* and *C. vagabundus* frequently and exclusively affiliate with their partners, displaying proximate swimming 75 and 81 % of the time, respectively; and paralleled swimming 53 and 69 % of the time, respectfully. Agonism in pairs is exclusively directed towards non-partner conspecifics, and is generally passive (i.e., dominated by visual or lateral displays and chasing is uncommon). Pairs exhibit strong partner fidelity, with 5/7 pairs of *C. baronessa* and 8/9 pairs of *C. vagabundus* maintaining their original partners throughout the duration of the study (6 weeks). Hence, we verify that (as in *C. lunulatus*)*, C. baronessa* and *C. vagabundus* are predominantly pair bonding at Lizard Island. Consistently, the predominant group size of *C. baronessa* and *C. vagabundus* is invariably paired across study populations. For *C. baronessa*, 70 % of observations at Heron Island (Australia) [32] and 55 % at Yaeyama Island (Japan) [74] are of paired groups. Whereas for *C. vagabundus*, 75 % of observation at Yaeyama Island (Japan) and Moorea Island (French Polynesia), and 65 % at Heron Island (Australia) populations are of paired groups. In both species, pairs maintain long-term territories that they defend against other butterflyfishes (*C. baronessa*: Heron Isl., Australia [32, 90]; *C. vagabundus*: Sesoko Isl., Japan [110]). However, previous descriptions of intra-pair relations are anecdotal, qualitative, and limited to one population of *C. baronessa* (Heron Island, Australia). Here, partners reportedly graze far apart within their territory, and only momentarily swim close together upon return from extra-territory forays [32]. For both species, pair sex composition has been previously examined in one population (Heron Isl., Australia), where all pairs are heterosexual [32]. Based on these aforementioned attributes of select populations, and despite further descriptions of partner relations or fidelity for *C. baronessa* or *C. vagabundus*, both species are nevertheless presumed to be pair bonding throughout their distributions [60]. While never explicitly tested or observed in *C. baronessa* or *C. vagabundus*, the prevalence of pair bonding does imply monogamous mating [5, 60]. However, spawning observations (as per [68]) are required for verification.

By contrast, we found that at Lizard Island, *C. trifascialis*, *C. rainfordi*, and *C. plebeius* all occur primarily as solitary individuals (80, 88, and 90 % of observations, respectively), and rarely in pairs (8-16% of observation) or aggregations (2-5 % of observations). Singletons exhibit no apparent social affiliation with another conspecific, as they spent 100 % of their time swimming further than 1.5 m from another conspecific; and most commonly, no other conspecific was within the field of view. Across its geographic range, *C. trifascialis* predominantly occurs as solitary individuals (Red Sea: 93 % of observations [58]; Moorea, French Polynesia: 86 % [73]; Heron Isl., Australia: 82 % [32]; Yaeyama Isl., Japan: 100 % [74]). Solitary grouping remains the predominant group size during the reproductive season, but not surprisingly, its prevalence can decline by 25 % (and pairing can increase by 25 %) (Yaeyama Isl., Japan) [74]. Adults establish long-term territories [90, 91], wherein territories of males’ encompass those of females’ [91]. In Kawashima (Japan), males repeatedly visit females within their territories, but spend only a short time swimming together (16 %) [91]. They moreover mate sequentially with inhabiting females, suggestive of haremic mating [91]. Singletons aggress against same sex conspecifics as a form of mate guarding [91] and against other butterflyfishes as territory defense [32, 101]. Social grouping of *C. rainfordi* has been previously examined at only Heron Isl. (Australia) [32]; where solitary individuals occur 98 % of the time. *Chaetodon pebeius* is also predominantly solitary at Heron Isl. (occurring 93 % of the time); yet, the species exhibits no predominant group size at Yaeyama Isl., (Japan), where it occurs equally as solitary and paired individuals (50 % of observations, respectfully), suggestive of variation in population-typical social system for the species. Solitary living in *C. ranifordi* and *C. plebeius* may be attributed to their more generalized diet [97], which conceivably reduces competition and consequently the need for cooperative territory defense. Although mating systems of *C. rainfordi* and *C. plebeius* are yet to be studied, they have been considered by some researchers to be monogamous [5]. However, the preponderance of solitary living and female-biased sex ratio found here suggests they are either polygynous or polygamous [60]. Clearly, more work is required to establish *C. rainfordi* and *C. plebeius* mating systems.

An unexpected and notable finding in this study was that one pair of *C. lunulatus* consisted of a female and of an individual simultaneously possessing both ovarian and testicular cells within their gonads. Gonads containing both sex cell types were also observed in one individual for each of the solitary species (*C. rainfordi, C. plebeius*, and *C. trifascialis*). A similar finding was previously reported once for chaetodontids, in a pairing and monogamous congener, *C. multicinctus*, who was histologically shown to occasionally exhibit spermatogenic tissue within ovaries [71]. These results tentatively suggest sequential hermaphroditism [111, 112] in these species, challenging the currently held view that chaetodontids are invariably gonochoric [113]. The additional observation of female-biased sex ratios in the 3 solitary species in this study further suggests protogynous hermaphroditism in in these species. These findings provide impetus for further substantiating sex change within these species and exploring its possible adaptive function(s) in relation to their social systems, e.g. [114, 115].

### Utility of study species for comparatively studying regulatory mechanisms of pair bonding: informing evolutionary history and controlling ecological confounds

Among the 6 study species, pair bonding is the ancestral state from which a single transition to solitary living likely occurred in the common ancestor of *C. rainfordi*, *C. plebeius*, and *C. trifascialis* (Fig 2). Such transitions from pair bonding to non-pair bonding systems are rare in animals, and represent a unique opportunity to serve as a “natural knock-out” for comparatively identifying pair bonding mechanisms, provided necessary controls are put in place.

Mechanisms that govern pair bonding independently of other behavioral or ecological factors remain poorly understood. This is partially because in most vertebrate groups, social systems naturally co-vary with these factors, making it very difficult to design highly controlled comparative systems [49]. Even in the most widely-used systems for comparatively studying pair bonding, *Microtus* voles and *Peromyscus* mice, species differences in pair bonding are confounded with species differences in parental care and habitat preference [8, 49]. This challenge persists in pre-existing teleost comparative systems. In the *Herichthys spp* cichlid system, pair bonding variation among males co-varies with variation in territoriality and parental care [48]. Whereas, in the teleost *Neolamprologus-Telmatochromis spp* system, species differences in pair bond strength are confounded with differences in group living and overall sociability [47, 52]. Finally, in a pre-established butterflyfish system, species differences in paired grouping *do not* co-vary with differences in other attributes (relatedness, aggression, or mating system) [31], and therefore are well controlled. However, because social group size does not fully characterise (and therefore nor does it verify) a social system [32, 60, 81, 87], the design may be invalid for comparative analysis of pair bonding specifically. Hence, additional, more controlled and validated systems for comparatively studying variation in pair bonding would be beneficial.

While we have established that the Lizard Island populations of the 6 study species display a clear variation in social systems, several key aspects of their behavioral ecology do not co-vary (Fig 5). All exist sympatrically at the location, where they are benthic feeders that (with the exception of *C. vagabundus*) feed almost exclusively on scleractinian corals [97, 99]; and exhibit differences in territoriality [101, 107] independently of differences in social system. These species, as in all butterflyfishes, are exclusively pelagic spawners, so are non-parental [116–118]. Importantly, all of these aforementioned ecological, behavioral, and geographic controls also apply for pair bonded vs. solitary individuals of *C. lunulatus* (Fig 5). Therefore, the proposed design offers a unique opportunity for highly controlled intra-and inter-species comparative research on the regulatory mechanisms of pair bonding. A logical next step would be to sample wild fish and compare mechanistic components (e.g., genes, neurochemicals, receptors, and brain regions) between pair bonding and solitary *C. lunulatus* individuals and/or *Chaetodon* species to identify mechanistic correlates of pair bonding.

**Fig 5.**
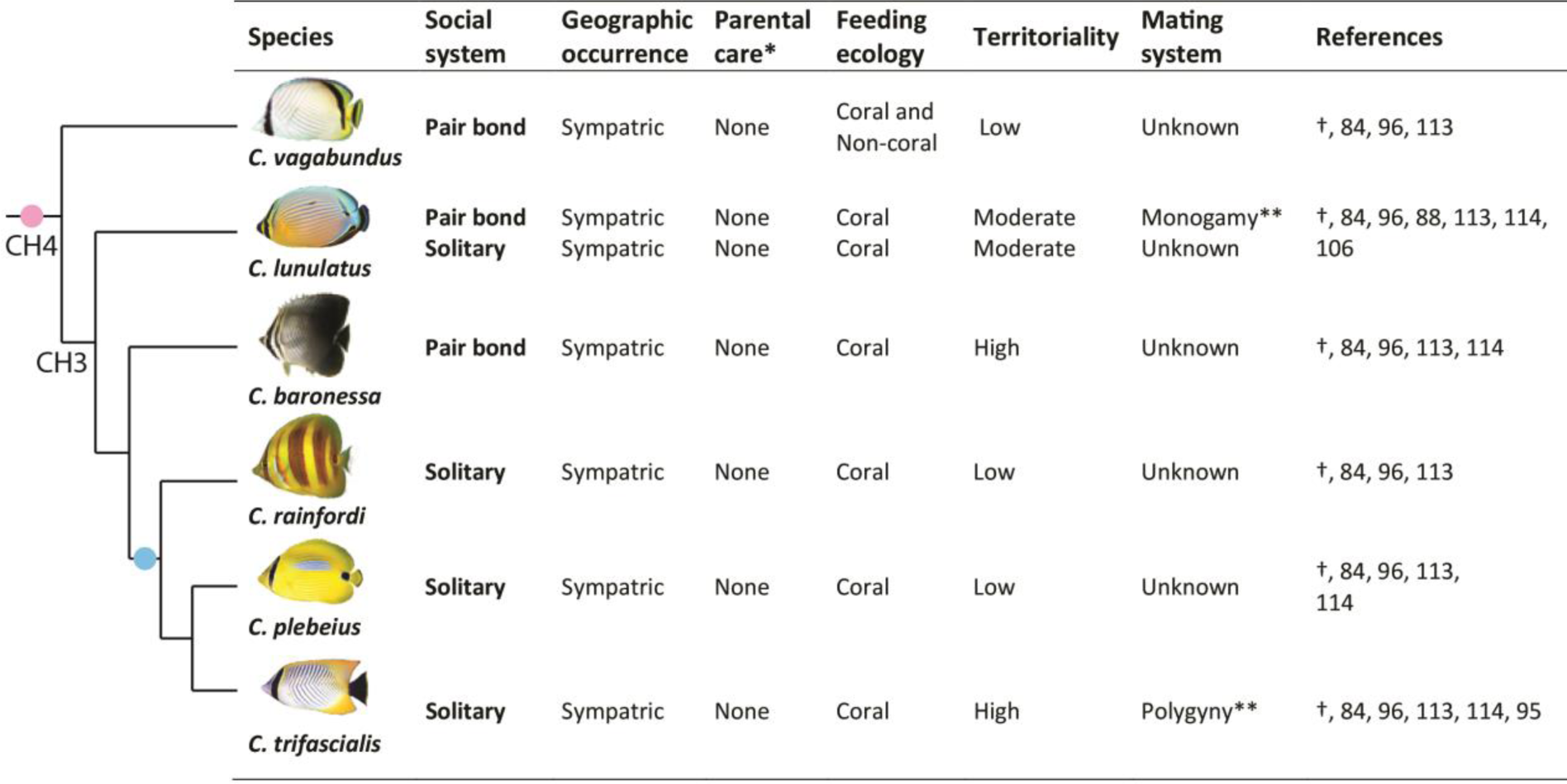
Dichotomous social systems (pair bonding vs. solitary living) among individuals of Chaetodon lunulatus and among species of Chaetodon at Lizard Island (current study) do not co-vary with other attributes (previously established), controlling for these variables while comparatively studying pair bonding. Phylogeny data from current study, where species clades are shown, and nodes represent species relatedness. Pink and blue circles indicate when pair bonding and solitary living evolved, respectfully based on stochastic character mapping. Notes: † = current study. *Parental care is unstudied in Lizard Island populations and is presumed absent based on unequivocal reporting of pelagic spawning within Chaetodontidae. **Mating systems of these populations at Lizard Island are presumed based on reports at other locations [68, 91].

Although the proposed *Chaetodon* butterflyfish system exhibits several attractive features for comparatively studying pair bonding, it does entail various limitations and challenges. As with most wild chaetodontids, most of these species have dietary requirements that are highly specialized and reliant on coral (*C. vagabundus* notwithstanding), making them difficult to maintain in captivity without growing coral; or changing their diet to one that is more economical/accessible (e.g., crustaceans, mussels, *Aiptasia* spp); which, although reportedly challenging, is achievable for even the most specialized corallivores, including *C. lunulatus* and *C. trifascialis* [119, 120]. While captive breeding of butterflyfish has been unsuccessful to date, it is expected to be achieved within the near future [119]. Until then, studies must be restricted to wild populations. Although these species are widely distributed relatively common, we cannot be certain that our findings on their social systems at Lizard Island translate to all populations/geographic locations. However, available data on their predominant group size and social behavior is highly consistent across populations/geographic locations (*C. plebeius* notwithstanding), indicating their social systems may be as well (*C. plebeius* notwithstanding once more). Verifying this should be a priority.

## Conclusions

In summary, this is the first study to examine the evolutionary history of Chaetodontidae sociality, revealing that within the family, pair bonding is ancestral and moderately conserved. It moreover verifies among 6 *Chaetodon* species at Lizard Island, Australia, a strong dichotomy in social systems representing one transition between them: from pair bonding in *C. lunulatus, C. baronessa, C. vagabundus* to solitary living in *C. trifascialis, C. rainfordi, C. plebeius*. These differences in social systems are not confounded with other life-history attributes. Therefore, these populations are useful for conducting controlled comparative analyses on the mechanistic correlates of pair bonding within an evolutionarily informed framework. A comparison of underpinning biological mechanisms found within the group to those in other emerging/established teleost, avian, and mammalian systems (among whom pair bonding has evolved independently), will help illuminate both general and dissociable mechanisms of pair bonding within vertebrates.

## Acknowledgements

We thank the anonymous referees for their thoughtful and constructive comments on earlier drafts of this manuscript. Thank you Andrew Cole, Marian Wong, Kelly Boyle, and Tim Tricas for logistic advice on butterflyfish capture and sexing. Manuela Giammusso, Kyvely Vlahakis, and Siobhan Heatwole provided excellent field assistance. David Hallmark kindly donated tagging equipment for this study. We thank Lizard Island Research Station for field support. We acknowledge all of the fishes that were sacrificed in order to undertake this project. This project was financially supported by ARC grants to M.S.P and S.P.W, and a GRS postgraduate research grant to J.P.N. Great Barrier Reef Marine Park Authority permit: G10/33239.1, G13/35909.1, G14/37213.1; James Cook University General Fisheries permit: 170251; James Cook University Animal Ethics approval: A1874.

## Supporting information

**S1 Table.** Species-typical social and mating systems in butterflyfishes.

